# The monosialoganglioside GM1a protects trophoblasts, erythrocytes and endothelial cells against complement attack

**DOI:** 10.1101/2022.01.24.477475

**Authors:** Henri Wedekind, Julia Beimdiek, Charlotte Rossdam, Elina Kats, Vanessa Wittek, Lisa Schumann, Andreas Tiede, Falk F. R. Buettner, Birgit Weinhold, Anja Münster-Kühnel, Markus Abeln

## Abstract

The complement system is a part of the innate immune system in the fluid phase and efficiently eliminates pathogens. However, its activation requires tight regulation on the host cell surface in order not to compromise cellular viability. Previously, we showed that loss of placental cell surface sialylation in mice *in vivo* leads to a maternal complement attack at the fetal-maternal interface, ultimately resulting in loss of pregnancy. To gain insight into the regulatory function of sialylation in complement activation, we here generated trophoblast stem cells devoid of sialylation, which also revealed complement sensitivity and cell death in vitro. Glycolipid-analysis by xCGE-LIF allowed us to identify the monosialoganglioside GM1a as a key element of cell surface complement regulation. Exogenously administered GM1a integrated into the plasma membrane of trophoblasts, dramatically increased binding of complement factor H and was sufficient to protect the cells from complement attack and cell death. Furthermore, GM1a treatment rescued sensitized human erythrocytes and endothelial cells from complement attack in a concentration dependent manner. This study demonstrates for the first time the complement regulatory potential of exogenously administered gangliosides and paves the way for sialoglycotherapeutics as a novel substance class for membrane-targeted complement regulators.

**Key points:**

- The naturally occurring sialic acid containing glycosphingolipid GM1a is a potent regulator of complement activation on mammalian trophoblast, erythrocyte and endothelial cell surfaces.
- Incorporated GM1a recruits complement factor H and thus could represent a novel membrane-targeted complement therapeutic.

## Introduction

The complement system is part of the innate immune system in the fluid phase and one of the first defense lines against pathogens. One mechanism by which complement activation confers protection against pathogens is the formation of pores in the cell surface, called membrane attack complex (MAC). While complement-mediated lysis represents an effective way to eliminate pathogenic entities, host cells require strict complement regulation to remain unscathed. All cells in the circulation are permanently exposed to low levels of complement activation. This includes circulating cells as well as endothelial cells and placental cells facing the maternal fluid phase. Excessive complement activation compromises host cell viability and is involved in numerous pathologic conditions.^1^

Immune homeostasis at the fetal-maternal interface is particularly intricate. The placental surface facing the maternal blood represents an interface between two genetically distinct entities and needs to establish immune tolerance to the embryo, while simultaneously ensuring potent immune defense against pathogens. This task is especially challenging in placentas of humans and rodents, in which the fetal trophoblast cells are in direct contact with the fluid phase of the maternal circulation. Loss of fetal-maternal immune balance results in pregnancy complications such as preeclampsia, or in severe cases, premature termination of pregnancy. Dysregulated complement activation was identified as a common pathway of injury in different human pregnancy complications.^2–4^ Recently we made use of a sialylation deficient mouse model and showed that sialylation plays a crucial role in placental complement regulation.^5^ Complete loss of fetal sialoglycans resulted in activation of the alternative pathway of the maternal complement system and ultimately embryonic death.

Loss of sialic acid on the cell surface can naturally occur due to its chemical instability, e.g. during aging or storage of cells for transfusion, or by the action of neuraminidases of endogenous or pathogenic origin.^6–8^ The influence of sialoglycans on complement regulation *per se* has been known for a long time and can mainly be attributed to the interaction of the complement regulatory protein factor H with cell surface sialoglycans. Factor H regulates complement activation by limiting the activation of the central complement component C3 to C3b by dismantling C3b generating C3 convertases and the recruitment of factor I, which proteolytically inactivates C3b to iC3b. Analysis of the interaction between sialoglycans and factor H culminated in the crystal structure of the factor H domains 19-20 with 3’sialyllactose (Neu5Acα2-6Galβ1-4Glc).^9^ Although the minimal binding motif for the sialoglycan-factor H interaction has been identified, it remained elusive whether sialoglycoconjugates could be exploited as complement regulatory reagents.

Here, we generated sialylation negative trophoblast stem cell lines, which were shown to be highly complement sensitive *in vitro*. Incorporation of the exogenously administered ganglioside GM1a into the membrane enhanced recruitment of the complement regulatory protein factor H to murine trophoblasts. Moreover, GM1a restored complement regulation also of human erythrocytes and endothelial cells. Thus, GM1a and other related gangliosides might represent a novel membrane resident complement regulator that could be used for future therapeutic intervention in disorders resulting from excessive complement activation.

## Methods

### Stem Cell Culture

The TS-Rs26 cell line, originally from the Rossant Lab (Toronto, Canada), was a kind gift of the Hemberger lab, Cambridge, United Kingdom. Standard culture conditions are described in Supplementary Methods. Transfection was performed using Lipofectamine (ThermoFisher) according to the manufacturer’s protocol. For treatment with sialyltransferase inhibitor, 3F_ax_-Peracetyl-Neu5Ac (Sigma-Aldrich) was added to a final concentration of 200 µM to the medium. Depletion of C3 from fetal bovine serum was performed using 0,83 µg/ml CVF (Quidel) for 30 min at 37 °C. Neuraminidase treatment of cells was conducted with 0,13 mg/ml purified *Arthrobacter ureafaciens* neuraminidase (in house production according to ^10^) in serum-free TSC medium for 30 min at 37 °C.

### Generation of *Cmas*^*-/-*^TSC

CRISPR-Cas9-mediated gene ablation was carried out using a guide RNA targeting *Cmas* exon 4 (5’-TGTCGACGAGGCCGTTTCGC-3’) in the plasmid pX330-U6-Chimeric_BB-CBh-hSpCas9 (Addgene plasmid #42230 from Feng Zhang) as used before.^11^ TSC were cotransfected with GFP-expressing plasmid peGFP-CI (Clontech) and sorted for GFP-positive cells in a 96 well plate at the central research facility cell sorting at MHH. Control TSC were mock transfected and also sorted. Screening for *Cmas*^*-/-*^ TSC was performed using primer targeting exon 4 (5’-GTCTGCATTCTGAGGGGAGT-3’ and 5’-AGAGCACAACACAGAAGGCT-3’) followed by Sanger sequencing (Eurofins).

### xCGE-LIF

As described previously, glycosphingolipid (GSL) glycosylation was assessed by multiplexed capillary gel electrophoresis coupled to laser induced fluorescence (xCGE-LIF) detection using an ABI PRISM® 3100-Avant Genetic Analyzer (advanced biolab service GmbH, Munich, Germany).^12^ The procedure is described more in detail in Supplementary Methods. Annotation of peaks was performed by migration time alignment to our in-house database.^12^

### qRT-PCR

RNA preparation with TRIzol® (invitrogen) and cDNA synthesis were performed according to the manufacturers protocols. Sequences of qPCR Primer (Sigma-Aldrich) are listed in Table S1. qPCRBIO SyGreen Lo-ROX Mastermix (Nippon Genetics Europe, Germany) was used on a QuantStudio 3 thermocycler. Ct values were normalized to three reference genes (Pgk1, Sdha, Ywhaz). Three biological replicates for each independent clone were used for all qPCR data.

### Immunofluorescence staining

For staining of cultured cells, TSC were seeded on glass coverslips. Fixation was performed in 4 % PFA/PBS and when needed permeabilised using 0.2 % Triton-X 100 for 10 min. Blocking and antibody incubations were carried out in 1 % BSA/PBS for 1 hour at RT. Detailed dilutions of used antibodies are listed in Table S2. Processed coverslips were mounted in Vectashield mounting medium (Vectorlabs, Burlingame, CA, USA), containing DAPI as counter-stain for nuclei. Images were taken with Zeiss Observer.Z1 microscope equipped with an AxioCam MRm digital camera and ApoTome module. Analysis was performed using Zeiss Zen software.

### Proliferation Assay

TSC were seeded in 96-well-plates at a density of 300 cells/well and incubated under differentiation conditions. Proliferation of differentiating TSC was analyzed using WST-1 reagent (Roche, Basel, Switzerland) 24 hours, 48 hours, 72 hours and 92 hours after seeding. 10 µl WST-1 reagent was added to the well and incubated for 4 hours at 37 °C. Measurement of absorbance at 450 nm and 690 nm as reference wavelength determined cell proliferation. This assay was performed three times with technical triplicates.

### GM1a incorporation of TSC

Prior to the incorporation of GM1a into the cell membrane, TSC were seeded in a 12-well plate and incubated at stem cell conditions. After 72 h, TSC were incubated for three hours with serum-free TSC medium and a concentration of 800 µM GM1a (Carbosynth Biosynth, United Kingdom) at 37°C. Subsequently they were either directly fixed as described earlier, or passaged to a new 12-well plate for differentiation.

### Complement Factor H binding assay

TSC were seeded in a 12-well plate and incubated at stem cell conditions for approximately 72 hours. GM1a incorporation was performed in selected wells as described above. Afterwards, TSC were washed two times with serum-free TSC medium. Incubation with human complement factor H (Complement Technology, Texas, USA) was carried out at 37°C for 30 min in serum-free TSC medium at a final concentration of 10 µg/ml. Subsequently, TSCs were directly fixed as described above and used for indirect immunofluorescence analysis.

### Western Blotting

Protein lysates were prepared in RIPA buffer and BCA Assay (Thermo Scientific) was used to determine protein concentration. Equal protein amounts were separated in a 12% SDS-PAGE and blotted onto PVDF membrane. Primary antibody incubation was performed over night at 4 °C with subsequent secondary antibody incubation for 1 hour at RT. Detailed dilutions of used antibodies are listed in Table S2. Enhanced chemoluminescence was utilised for detection.

### Hemolysis assay

The hemolysis assays using human and sheep red blood cells are described in detail in the Supplementary Methods.

### Calcein release assay

The assay was performed as described before.^13^ Description of the procedure can be found in Supplementary Methods.

### Statistical Analysis

Statistical analysis was performed using GraphPad Prism 7 software (GraphPad, San Diego, CA, USA). One-way ANOVA and mixed two-way ANOVA followed by Bonferroni’s post hoc test were applied as indicated. Normality and equality of variances were assessed by the Shapiro–Wilk and the Brown–Forsythe test, respectively. * P < 0.05, ** P < 0.01, *** P < 0.001.

## Results

### Generation of asialo trophoblast stem cells

Genetic inactivation of the *Cmas* gene in the murine trophoblast stem cell (TSC) line TS-Rs26 was achieved by applying the CRISPR/Cas9 technology (**Fig. 1A**). The guide RNA was designed to target the active site of the *Cmas* gene (**Fig. S1A**,**B**). Clonal lineages were established and clones with frame-shifts in both alleles were viable and termed *Cmas*^*-/-*^ (**Fig. S1B**,**C**). Three *Cmas*^*-/-*^ TSC lines were used for all further analyses. Loss of CMAS expression in *Cmas*^*-/-*^ TSC lines was confirmed by Western Blot analysis (**Fig. 1B**). While control cells showed a signal at the expected molecular weight at 48 kDa, all three *Cmas*^*-/-*^ TSC lines lost the corresponding CMAS signal (**Fig. 1B**). Evaluation of different characteristic TSC markers by qPCR and immunofluorescence did not reveal major differences, demonstrating that loss of CMAS did not generally alter the stem cell character of TSC (**Fig. S1D**,**E**).

**Figure 1.**
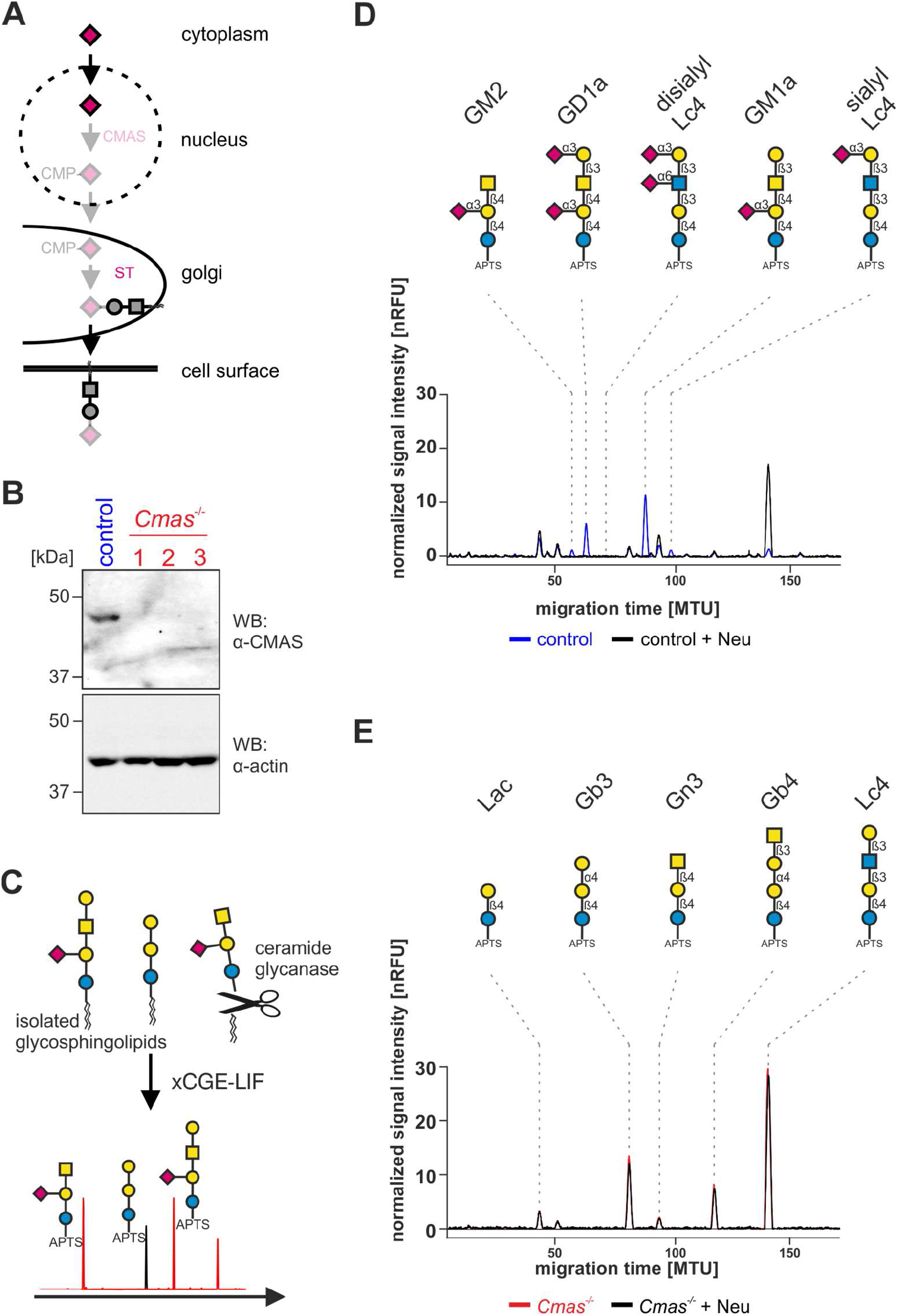
*Cmas*^*-/-*^trophoblast stem cells are devoid of sialoglycans. (**A**) Scheme of the biosynthesis of sialoglycoconjugates. Genetic inactivation of *Cmas* results in unsialylated glycans at the cell surface. (**B**) CMAS Western blot analysis. Control and *Cmas*^*-/-*^ TSC were lysed and separated by SDS-PAGE and immunostained with anti-CMAS antibody. Anti-actin immunostaining was used as loading control. (**C**) Scheme of xCGE-LIF of glycosphingolipids. The glycan head group is released by cleaveage with a ceramide glycanase. After labelling with the fluorescent dye APTS, glycans are separated using capillary gel electrophoresis depending on their physicochemical properties. (**D-E**) xCGE-LIF analysis of glycans derived from glycosphingolipids. Glycan notations for GM2, GD1a and GM1a refer to the respective glycosphingolipid derived glycan. Overlay of xCGE-LIF electropherograms of APTS-labeled GSL-derived glycans of control (**D**) and *Cmas*^*-/-*^ TSC (**E**) with or without prior treatment with Neu. One representative *Cmas*^*-/-*^ TSC clone is shown. For inter-sample comparisons signal intensities were normalized to Man6 (nRFU). MTU: migration time unit. Symbols nomenclature according to ref ^33^.

Depletion of CMAS in murine embryonic stem cells has led to a complete loss of cell surface sialic acid ^14^. To exemplarily determine the sialylation status of control and *Cmas*^*-/-*^ TSC glycans, we used multiplexed capillary gel electrophoresis coupled to laser-induced fluorescence detection (xCGE-LIF) to analyse glycosphingolipid glycosylation (**Fig. 1C**). Glycosphingolipid derived glycans of control cells were analysed with and without prior enzymatic cleavage of sialic acid residues by Neuraminidase (Neu) from *Arthrobacter ureafaciens*. xCGE-LIF demonstrated that control TSC possessed two major (GD1a and GM1a) and three minor ganglioside species (GM2, disialyl-Lc4-Cer and sialyl-Lc4-Cer) (**Fig. 1D**). *Cmas*^*-/-*^ TSC however entirely lacked all ganglioside species, but exhibited markedly elevated levels of the neutral glycosphingolipids lactosylceramide (Lac-Cer), globotriaosylceramide (Gb3-Cer), gangliotriaosylceramide (Gn3-Cer), globotetraosylceramide (Gb4-Cer) and lactotetrasylceramide (Lc4-Cer) (**Fig. 1E, S2A**,**B**). Hence, *Cmas*^*-/-*^ TSC globally lack sialylated structures on glycosphingolipids and mutant cells are further called “asialo” (**Fig. 1A**).

### Asialo trophoblast stem cells are sensitive to complement activation

Control and *Cmas*^*-/-*^ TSC were analyzed regarding their differentiation capacities by cultivation in differentiation medium. In a WST-1 cell proliferation assay, control TSC showed increasing proliferation over the whole differentiation time period of 96 h, while *Cmas*^*-/-*^ TSC revealed a marked reduction in proliferation (**Fig. 2A**). Based on our previous observations *in vivo* ^5^ the rapid decrease of *Cmas*^*-/-*^ TSC cell viability upon induction of differentiation made the contribution of immune-active factors in the differentiation medium likely. To analyze whether differentiating *Cmas*^*-/-*^ TSC activate the complement system *in vitro*, we examined the presence of the central complement component C3 by indirect immunofluorescence. As a control to complement active serum, heat inactivation of serum or treatment with cobra venom factor (CVF) deprived the serum in the medium of active components of the complement cascade (**Fig. S3**). While control TSC showed no reactivity, *Cmas*^*-/-*^ TSC revealed increased deposition of C3 on the cell surface (**Fig. 2B**), which was abolished when heat-inactivated or CVF treated serum was used (**Fig. 2B**). These results clearly demonstrate that loss of TSC viability is accompanied by activation of the complement system.

**Figure 2.**
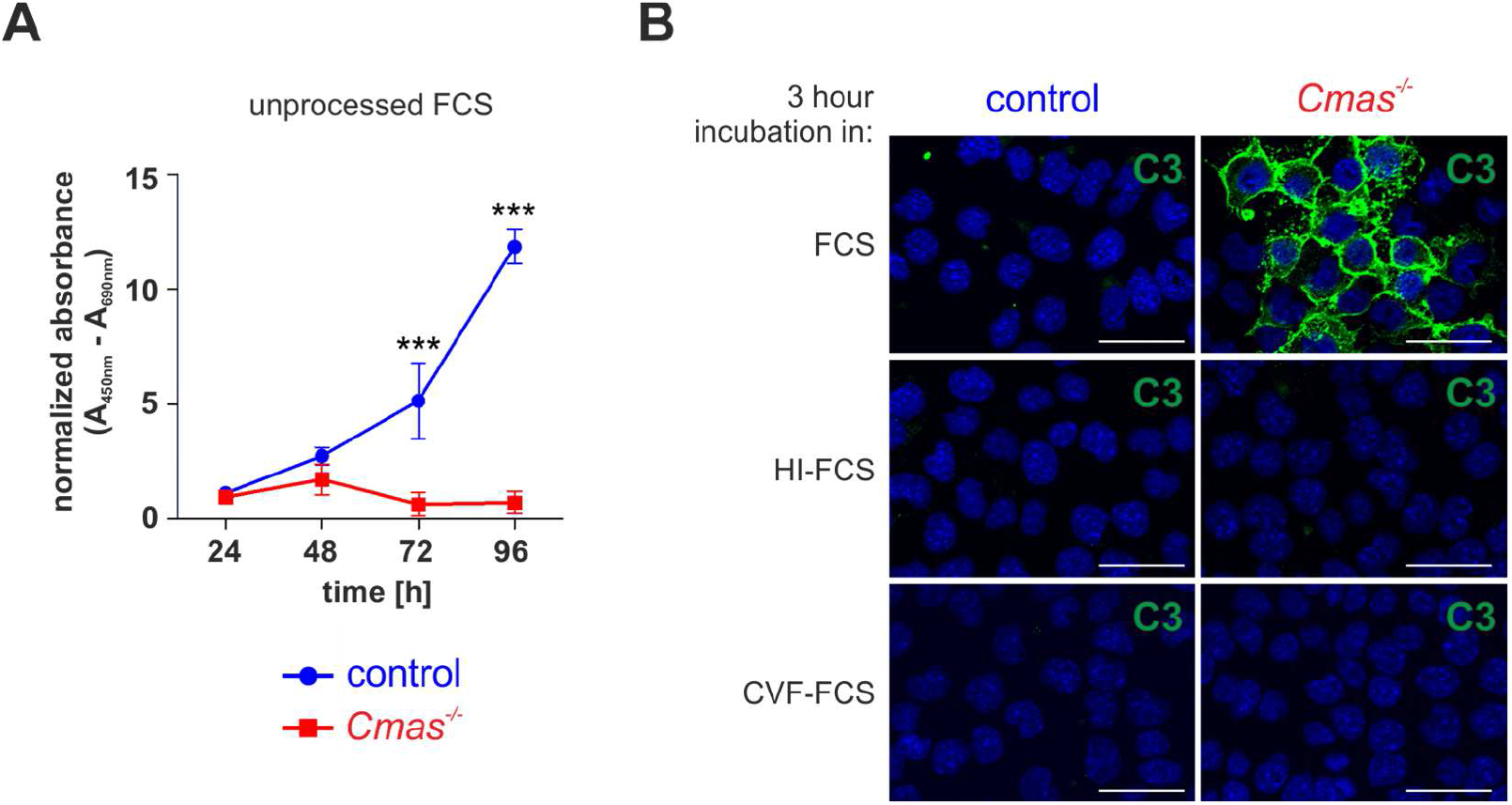
*Cmas*^*-/-*^TSC are only able to differentiate in heat inactivated serum. (**A**) WST-1 proliferation assay of control and *Cmas*^*-/-*^ TSC in differentiation medium with unprocessed FCS. Absorbance was measures every 24 hours after seeding for four days in three independent experiments. For statistical analysis a two-way ANOVA with Bonferroni’s multiple comparison post-test was performed *** P < 0.001. Mean values are depicted with standard deviation. (**B**) Detection of C3 deposition by indirect immunofluorescence analysis of control and *Cmas*^*-/-*^ TSC in medium with untreated, heat inactivated (HI-FCS) or CVF treated (CVF-FCS) FCS for 3 hours. Nuclei were stained with DAPI and are shown in blue. Scale bars = 50 µm. Data from one representative *Cmas*^*-/-*^ TSC clone are shown.

Since differentiation of *Cmas*^*-/-*^ TSC was possible under complement deprived conditions, we subsequently examined whether loss of Sia has an impact on the differentiation capacity of TSC by quantifying the expression of genes associated with TSC stemness and differentiation. qPCR analyses revealed no differences between control and *Cmas*^*-/-*^ TSC (**Fig. S4A**,**B**). Control and *Cmas*^*-/-*^ TSC similarly downregulated the stemness associated genes *Elf5* and *Esrrb* and at the same time upregulated the differentiation markers *Prl3d1* and *Tpbpa* after 7 days of differentiation.

To verify that the deletion of *Cmas* was the primary reason for the observed complement sensitivity and to exclude off target effects, *Cmas*^*-/-*^ TSC were transfected with murine *Cmas*. Complementation of *Cmas*^*-/-*^ TSC not only restored sialylation (**Fig. S5A**,**B**), but also viability and differentiation in complement-active medium, demonstrating that loss of CMAS and subsequent loss of cell surface sialylation was responsible for the demise of *Cmas*^*-/-*^ TSC during differentiation (**Fig. S5C**).

### GM1a and sialyl-Lc4-Cer sialylation are sufficient to suppress complement activation

To confirm that the observed complement sensitivity results from the absence of cell surface sialoglycans, we made use of an approach that does not genetically, but chemically interferes with sialoglycan biosynthesis. The glycomimetic 3F-Neu5Ac interferes with sialylation of glycans by inhibition of the Golgi resident sialyltransferases (**Fig. 3A**).^15^ However, we observed that 3F-Neu5Ac treated control TSC showed no reactivity for C3 at the cell surface (**Fig. 3B**) and thus were less complement-sensitive than *Cmas*^*-/-*^ TSC. To answer whether the presence of residual sialoglycan species might be responsible for this observation, glycans derived from 3F-Neu5Ac treated control TSC were subsequently treated with Neu, which again resulted in complement sensitivity and C3 deposition (**Fig. 3B**). These results indicated that loss of sialylation by 3F-Neu5Ac treatment was indeed incomplete and the remaining sialoglycans might possess complement regulatory functions on the cell surface. We used xCGE-LIF analysis of glycosphingolipids from 3F-Neu5Ac treated control TSC to directly demonstrate residual sialylation. 3F-Neu5Ac treated TSC exhibited two remaining sialylated glycosphingolipids (**Fig. 3C,D**) identified as the GM1a derived glycan and sialyl-Lc4 by migration time alignment.^12^

**Figure 3.**
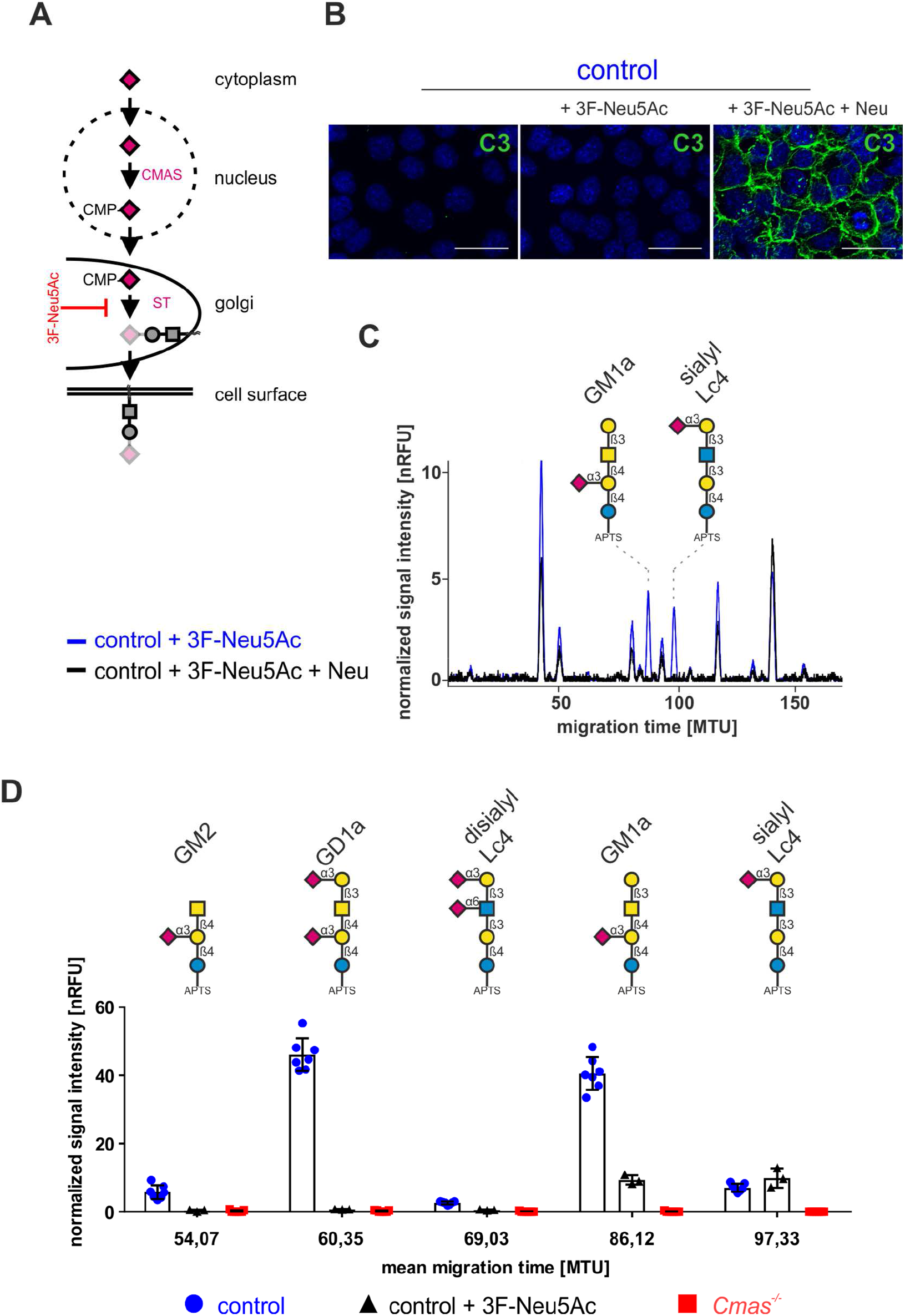
3F-Neu5Ac treated TSC are not devoid of gangliosides. (**A**) Scheme of the inhibition of sialyltransferases in the biosynthesis of sialoglycoconjugates by the glycomimetic 3F-Neu5Ac. (**B**) Detection of C3 deposition by indirect immunofluorescence analysis of untreated or 3F-Neu5Ac treated control TSC with and without subsequent Neu treatment. Nuclei were stained with DAPI and are shown in blue. Scale bars = 50 µm. (**C**) Overlay of xCGE-LIF electropherograms of APTS-labeled GSL-derived glycans of 3F-Neu5Ac treated control TSC with or without prior treatment with Neu. Glycan notation for GM1a refers to the glycosphingolipid derived glycan. For inter-sample comparisons signal intensities were normalized to Man6 (nRFU). MTU: migration time unit. Symbol nomenclature according to ^33^. (**D**) Quantification of GSL-derived glycans in untreated and 3F-Neu5Ac treated control TSC, and *Cmas*^*-/-*^ TSC using xCGE-LIF. Glycan notations for GM2, GD1a and GM1a refer to the respective glycosphingolipid derived glycan. Cut off was set to 2 nRFU. n=3 for control+3F-Neu5Ac; n=7 for control and *Cmas*^*-/-*^. Mean values are depicted with standard deviation.

### GM1a rescues regulation of complement activation on asialo trophoblast stem cells

This led us to the question whether GM1a and sialyl-Lc4-Cer directly contributed to the observed differences in TSC viability. As sialyl-Lc4-Cer is neither well studied nor commercially available and GM1a is known to incorporate into the plasma membrane *in vivo* and *in vitro* ^16,17^, we focused on GM1a. Although previous studies showed that GM1a exerts complement regulatory functions in artificial liposomes ^18^, other data suggests that ganglioside incorporation might even increase the susceptibility for complement mediated lysis.^19^ To clarify this issue, we assessed whether GM1a is able to regulate complement activation in *Cmas*^*-/-*^ TSC. GM1a was administered to control as well as to *Cmas*^*-/-*^ TSC cultures for 3 hours and successful GM1a incorporation into the plasma membrane was controlled by xCGE-LIF (**Fig. S6A**,**B**) and binding of the GM1a ligand Cholera toxin subunit B (CtxB). The vast majority of CtxB reactivity of control TSC was confined to the cell surface (**Fig. 4A**). However, cell surface reactivity of CtxB was completely abolished in *Cmas*^*-/-*^ TSC, but could be restored upon treatment with GM1a (**Fig. 4A**). Therefore, TSC are generally capable of incorporating GM1a into their plasma membrane.

**Figure 4.**
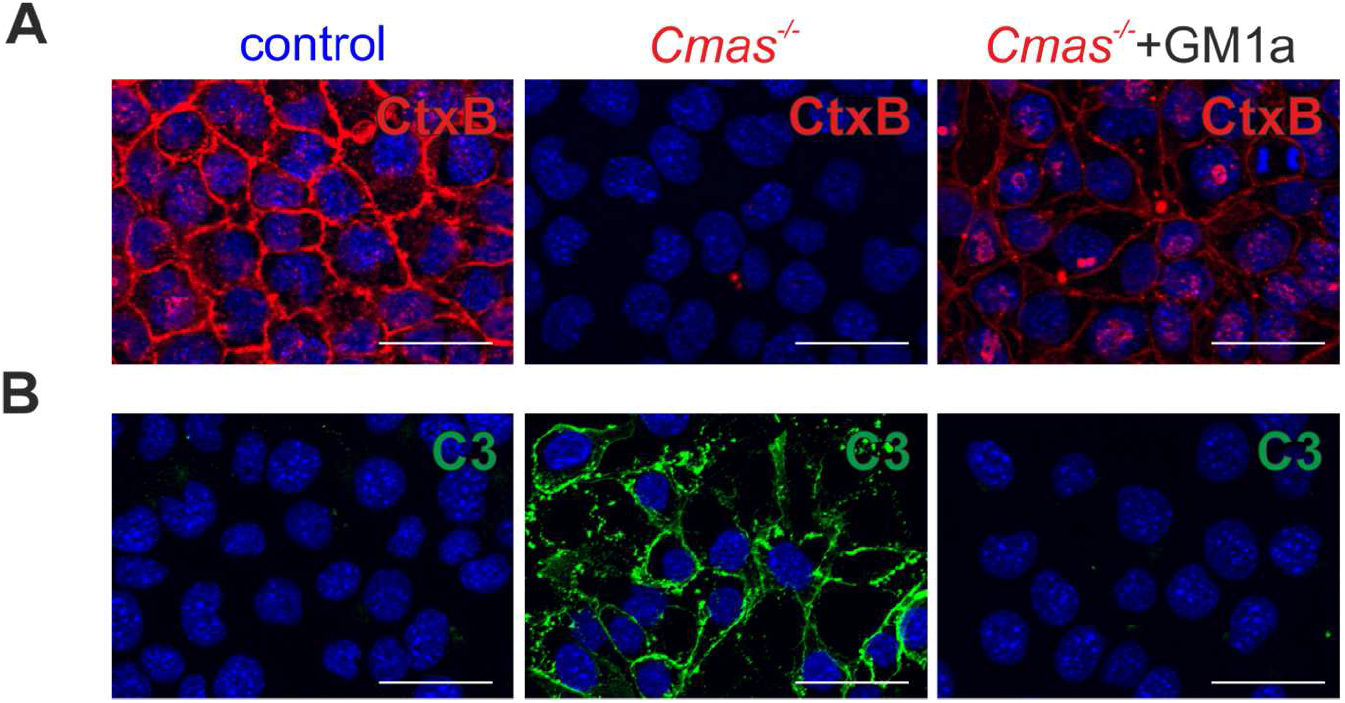
Exogenously administered GM1a protects *Cmas*^*-/-*^TSC from complement attack. (**A**) Detection of endogenous and incorporated GM1a with Cholera toxin subunit B (CtxB) in control TSC, *Cmas*^*-/-*^ TSC and *Cmas*^*-/-*^ TSC treated with GM1a in medium with untreated FCS. Scale bars = 50 µm. (**B**) Detection of C3 deposition by indirect immunofluorescence analysis of control, *Cmas*^*-/-*^ TSC and *Cmas*^*-/-*^ TSC treated with GM1a in medium with untreated FCS. Nuclei were stained with DAPI and are shown in blue. Scale bars = 50 µm. For all experiments is n=3 and were performed with all three *Cmas*^*-/-*^ TSC clones. One representative *Cmas*^*-/-*^ TSC clone is shown.

To answer whether GM1a incorporation is sufficient to rescue complement defense on *Cmas*^*-/-*^ TSC in standard differentiation medium, *Cmas*^*-/-*^ TSC were incubated with GM1a for 3 hours immediately before differentiation. Analysis of C3 deposition revealed that GM1a treatment markedly decreased the C3 positive population, demonstrating that GM1a incorporation improved the ability of *Cmas*^*-/-*^ TSC to regulate complement activation on the cell surface (**Fig. 4B**).

### GM1a exerts complement regulatory functions on erythrocytes and endothelial cells

The potent complement regulatory properties of GM1a in *Cmas*^*-/-*^ TSC raised the question if this feature of GM1a was applicable to other complement sensitive biological systems. Therefore, we analyzed whether GM1a is also capable of regulating human complement on sensitized human red blood cells (hRBC). hRBC were sensitized for complement mediated lysis by a functional blockade of the complement regulatory protein CD59 with a monoclonal antibody.^13^ To analyze hRBC with deficits in regulation of complement attack initiation we subjected hRBC to Neu treatment with or without subsequent GM1a supplementation and compared them to untreated hRBC. Control hRBC showed lysis of about 27 % that increased to 41 % after Neu treatment. Administration of GM1a to Neu-treated hRBC again decreased lysis to 27 % (**Fig. 5A**). While Neu treatment significantly sensitized hRBC to complement mediated lysis, GM1a treatment markedly reduced complement induced lysis similar to control levels. Another frequently used cellular system to assess complement regulatory potential of novel therapeutic approaches are Neu treated sheep red blood cells (SRBC), which are known to show complement dependent lysis if confronted with human serum and thus represent a well-studied model of complement activation.^20^ We confirmed the restoration of complement resistance by GM1a administration observed in hRBC also in SRBC (**Fig. S7A**,**B**). Summarized, exogenously applied GM1a was incorporated by human and sheep RBC and exerted complement regulatory functions.

**Figure 5.**
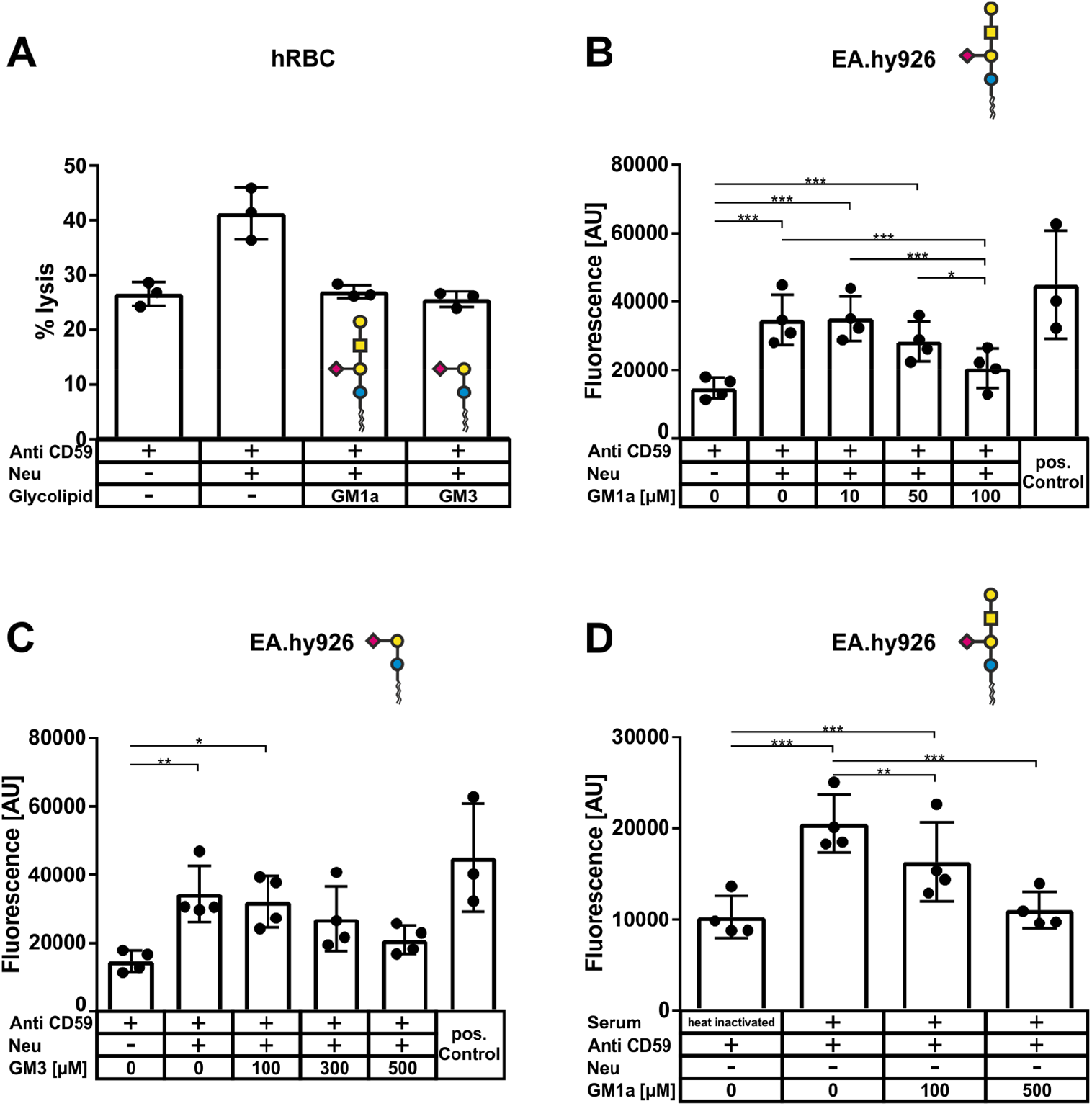
GM1a and GM3 protect against human complement. (**A**) Control, Neu sensitized or Neu sensitized and subsequently GM1a or GM3 (both 50 µM) treated human RBC were incubated with human complement active serum in DGHB-Mg-EGTA. Hemolysis was plotted relative to human RBC incubated with only red cell lysis buffer, which was defined as 100% lysis. All cells were treated with anti-CD59 to block the inhibition of MAC formation. (**B**) GM1a improves complement regulation on the endothelial cell line EA.hy926. A Calcein release assay was used to determine the complement activation on the human endothelial cell line EA.hy926. Cells were desialylated with Neu and subsequently incubated with different concentrations of GM1a. After treatment with human serum, the supernatants were measured for fluorescence at 494 nm. Shown are absolute fluorescent values and mean values ±SD. All cells were treated with anti-CD59 to block the inhibition of MAC formation. For the positive control, cells were lysed with detergents to release the entire Calcein. (**C**) GM3 improves complement regulation on the endothelial cell line EA.hy926. The same calcein release assay as in B was performed with GM3 instead of GM1a treatment. (**D**) GM1a improves complement regulation on the endothelial cell line EA.hy926 also without prior removal of sialic acid. The same calcein release assay as in (B) was performed but without neuraminidase treatment. For statistical analysis a repeated measures ANOVA with Bonferroni’s multiple comparison post-test was performed * P < 0.05, ** P < 0.01, *** P < 0.001. Mean values are depicted with standard deviation.

Another cell type that is prone for complement mediated injury is the endothelium, which *in vivo* is permanently exposed to complement proteins. To examine whether GM1a is also capable of regulating complement mediated injury on endothelial cells we used the human endothelial cell line EA.hy926 in a calcein release assay that was previously used to study complement activation on endothelial cells.^13^ Control cells without further treatment showed a mean fluorescent signal of 14702 ± 3069 AU in the cell culture supernatant after exposure to complement active serum (**Fig. 5B**). Cells that were additionally treated with Neu had a significantly higher fluorescent signal of 34622 ± 7378 AU, indicating increased calcium release due to MAC complex formation as a result of complement activation. Additional exposure with 100 µM GM1a abolished this difference. Furthermore, treatment with 100 µM GM1a also showed significantly decreased signals compared to non or 10 µM GM1a treated cells. These data show that exogenously administered GM1a also regulated complement activation on complement sensitized endothelial cells. Next, we asked whether in addition to GM1a also related gangliosides with similar glycan moieties possess complement regulatory functions and tested as a proof of principle the monosialodihexosylganglioside GM3 in the calcium release assay (**Fig. 5C**). GM3 is the most common membrane-bound glycosphingolipid in tissues and milk ^21^ and is composed of the same sialic acid bearing core structure (Cer-Glc-Gal-Sia) as GM1a but lacks the additional extension with *N*-acetylgalactosamine and galactose (**Fig. 5A**). GM3 treatment of sensitized hRBC produced the same effect of reduction in complement induced lysis as GM1a (**Fig. 5A**). Moreover, GM3 was also able to protect neuraminidase sensitized EA.hy926 cells against complement activation, but higher concentrations compared to GM1a treatment were necessary to visualize the effect (500 µM GM3 vs. 100 µM GM1a) (**Fig. 5C**). Next, we analyzed whether the complement regulatory function of GM1a also holds true for non-desialylated endothelial cells. Treatment of EA.hy926 cells with 500 µM GM1a decreased fluorescence signals in the calcein release assay further to a level comparable to cells exposed to heat-inactivated serum (**Fig. 5D**). This indicates that the complement regulatory function of incorporated GM1a is not restricted to cells with impaired sialylation.

Since factor H plays a crucial role in complement regulation and is known to interact with sialylated glycoconjugates ^22^, we assessed recruitment of purified human factor H to the cell surface with and without prior administration of GM1a. Factor H was not detectable on untreated TSC and EA.hy926 cells. Incorporation of exogenous GM1 increased detection and therefore binding of human complement factor H on both cell lines (**Fig. 6**). This indicated that exogenously administered and incorporated GM1a represents a recognition motif for complement factor H on murine TSC and EA.hy926 cells and suggests the mechanism for the complement regulatory function of GM1a.

**Figure 6.**
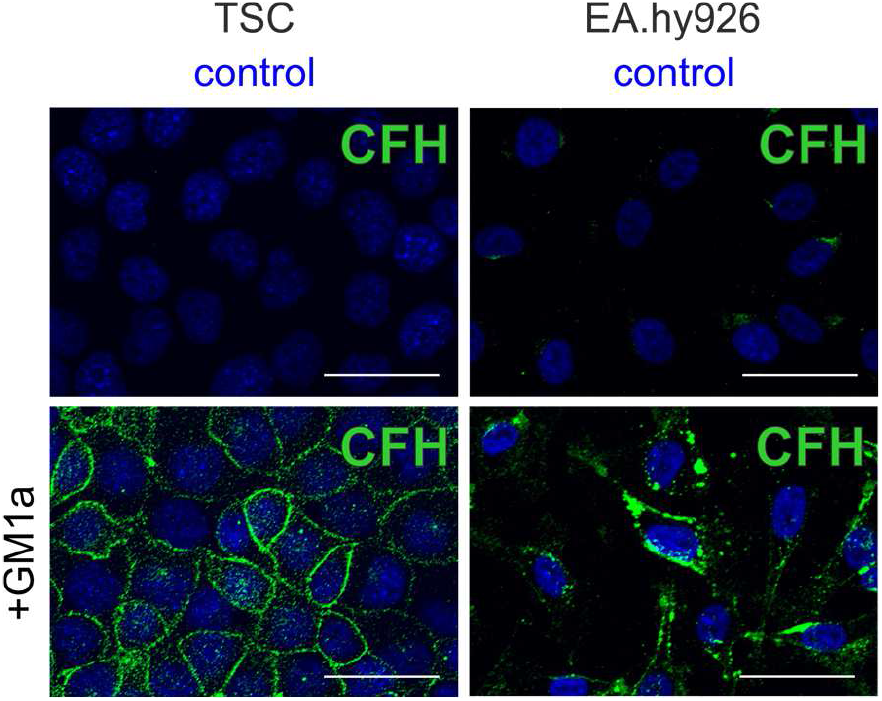
Complement factor H binds to GM1a. Detection of human complement factor H by indirect immunofluorescence analysis of control TSC and EA.hy926 with and without prior incubation with GM1a in medium with unprocessed FCS. Nuclei were stained with DAPI and are shown in blue. Scale bars = 50 μm.

## Discussion

The complement system requires tight regulation in order not to compromise host cell viability. Therefore it is not surprising that numerous pathologies include dysregulation of the complement pathway (e.g. paroxysmal nocturnal hemoglobinuria (PNH), C3 glomerulopathy, atypical hemolytic uremic syndrome (aHUS) and ischemic reperfusion injury).^1^ Most therapeutic approaches address exuberant complement activity by inhibiting C3 or C5 convertases, e.g. AMY-101 (Amyndas) a peptide inhibitor of C3 activation and blockage of C5 activation by C5 binding of the monoclonal antibody eculizumab. While these strategies effectively inhibit full-blown complement activation, they also systemically inhibit complement activation at the risk of infections with encapsulated bacteria and do not address the true underlying pathology of deficient complement defense on the host cell surface.^23,24^ Restoration of complement regulation directly on the cell surface would require complement defense molecules that are able to integrate into the plasma membrane of affected host cells. A proof that this concept is generally working is the membrane-targeted complement inhibitor Mirococept (APT070), which has been successfully employed in *in vivo* studies, including improved graft survival of cold stored donor organs in a rat transplantation model.^25^ However, production of Mirococept is complex and includes recombinant expression of SCR 1-3 of complement receptor type 1 in inclusion bodies, which then additionally need to be refolded and myristoylated *in vitro*.

Here, we describe the discovery of a novel function of the monosialoganglioside GM1a as a membrane targeted complement regulatory molecule. GM1a showed complement inhibiting potential as an endogenously present molecule or after integration into the cell surface when exogenously added as liposomes. Previous studies attributed to ganglioside treatment of hRBC an increase in complement sensitivity ^19^, while GM1a incorporation into artificial liposomes reduced complement activation.^18^ The complement regulatory potential of GM1a observed in our study expands these earlier observations and clearly demonstrates that GM1a integrates into the plasma membrane of hRBC, endothelial and placental cells and negatively regulates complement activation. Another important result of our study is that the complement regulatory potential of GM1a was not restricted to sensitized desialylated cells, as GM1a was also able to reduce complement mediated cellular damage in endothelial cells displaying native sialylation pattern. Furthermore, our studies revealed that exogenously added GM1a markedly increased recruitment of the complement regulatory protein factor H to the cell surface of control TSC and EA.hy926. Factor H consists of 20 complement-control protein (CCP) domains and is *inter alia* recruited to the cell surface via C3b binding sites; therefore, it preferably binds to cell surfaces with active complement events. Moreover, factor H possesses a Sia binding pocket that accommodates cell surface glycans with a terminal sialic acid in α-2,3 linkage and STD NMR analyses showed that factor H interacts with the glycan portion of GM1a and GM3.^9,26^ While recruitment of factor H could readily explain the complement regulatory potential of exogenous GM1a, it is not clear why factor H recruitment was not detected in control cells bearing endogenous GM1a. One explanation might be that GM1a treatment raised GM1a concentrations beyond endogenous levels. Additionally, it was shown that exogenous administration of GM1a induces changes in the organization of the plasma membrane, which could make GM1a and other factor H receptors more readily accessible.^27^ While we still lack a precise understanding of how different cell surface glycans contribute to the interaction of factor H with the cell surface, our data show that glycomic approaches like xCGE-LIF offer a great opportunity to identify glycans with such potential in future studies.

Our data emphasize once more the complement sensitive nature of trophoblast cell and provide first evidence that exogenously administered GM1a has complement regulatory functions in important cellular systems of the hematological compartment, including murine placental cells, human erythrocytes and human endothelial cells. GM1a is an intensely studied ganglioside that is produced naturally in the human body. It accounts for about 28% of the total human brain gangliosides and is currently being studied in clinical trails in the treatment of neurological diseases on the basis of its well-known neuroprotective function.^17,28,29^ Importantly, these clinical trials showed no adverse immuno-stimulant effects after injection of GM1a, suggesting that GM1a treatment as therapeutic intervention in complement related diseases would be similarly safe.^17,28^

Taken together our findings highlight the potential of GM1a, and possibly other gangliosides, as new therapeutic agents for membrane-targeted complement inhibition which might also be of interest for many different pathologies, including neurodegenerative, hemolytic, renal and autoimmune diseases.^1,30–32^

## Acknowledgments

We thank Myriam Hemberger (Calgary, Canada), Vicente Perez-Garcia (Cambridge, UK) and the Rossant lab (Toronto, Canada) for kindly providing the TS-Rs26 cell line, Mania Ackermann and Nico Lachmann (Hannover, Germany) for the EA.hy926 cell line. Also, we would like to acknowledge the assistance of the Cell Sorting Core Facility of the Hannover Medical School supported in part by Braukmann-Wittenberg-Herz-Stiftung and Deutsche Forschungsgemeinschaft. We thank Søren Christensen for providing the *Arthrobacter ureafaciens* sialidase expression plasmid and Rita Gerardy-Schahn for continuous support and inspiring discussions.

## Data Availability Statement

The datasets generated during and/or analyzed during the current study are available from the corresponding author on reasonable request.

## Authorship Contributions

H.W., J.B., C.R., E.K., V.W., L.S., E.K., and M.A. performed experiments; H.W., J.B., C.R., FFR.B., and M.A. analyzed and interpreted the results; H.W., J.B., and M.A. made the figures; M.A. and A.M.K. designed the research; H.W., B.W., A.T., A.M.K., and M.A. wrote the paper.

## References

1. Ricklin, D., Reis, E. S. & Lambris, J. D. Complement in disease: a defence system turning offensive. Nature reviews. Nephrology 12, 383–401; 10.1038/nrneph.2016.70 (2016).

2. Banadakoppa, M., Balakrishnan, M. & Yallampalli, C. Common variants of fetal and maternal complement genes in preeclampsia: pregnancy specific complotype. Scientific reports 10, 4811; 10.1038/s41598-020-60539-9 (2020).

3. Buurma, A. et al. Preeclampsia is characterized by placental complement dysregulation. Hypertension (Dallas, Tex. : 1979) 60, 1332–1337; 10.1161/HYPERTENSIONAHA.112.194324 (2012).

4. Hoffman, M. C., Rumer, K. K., Kramer, A., Lynch, A. M. & Winn, V. D. Maternal and fetal alternative complement pathway activation in early severe preeclampsia. American journal of reproductive immunology (New York, N.Y. : 1989) 71, 55–60; 10.1111/aji.12162 (2014).

5. Abeln, M. et al. Sialic acid is a critical fetal defense against maternal complement attack. The Journal of clinical investigation 129, 422–436; 10.1172/JCI99945 (2019).

6. Huang, Y.-X. et al. Human red blood cell aging: correlative changes in surface charge and cell properties. Journal of cellular and molecular medicine 15, 2634–2642; 10.1111/j.1582-4934.2011.01310.x (2011).

7. Li, M.-F. et al. Platelet desialylation is a novel mechanism and a therapeutic target in thrombocytopenia during sepsis: an open-label, multicenter, randomized controlled trial. Journal of hematology & oncology 10, 104; 10.1186/s13045-017-0476-1 (2017).

8. Soslau, G. & Giles, J. The loss of sialic acid and its prevention in stored human platelets. Thrombosis research 26, 443–455; 10.1016/0049-3848(82)90316-4 (1982).

9. Blaum, B. S. et al. Structural basis for sialic acid-mediated self-recognition by complement factor H. Nature chemical biology 11, 77–82; 10.1038/nchembio.1696 (2015).

10. Christensen, S. & Egebjerg, J. Cloning, expression and characterization of a sialidase gene from Arthrobacter ureafaciens. Biotechnology and applied biochemistry 41, 225–231; 10.1042/BA20040144 (2005).

11. Niculovic, K. M. et al. Podocyte-Specific Sialylation-Deficient Mice Serve as a Model for Human FSGS. Journal of the American Society of Nephrology : JASN 30, 1021–1035; 10.1681/ASN.2018090951 (2019).

12. Rossdam, C. et al. Approach for Profiling of Glycosphingolipid Glycosylation by Multiplexed Capillary Gel Electrophoresis Coupled to Laser-Induced Fluorescence Detection To Identify Cell-Surface Markers of Human Pluripotent Stem Cells and Derived Cardiomyocytes. Analytical chemistry 91, 6413–6418; 10.1021/acs.analchem.9b01114 (2019).

13. Hyvärinen, S., Meri, S. & Jokiranta, T. S. Disturbed sialic acid recognition on endothelial cells and platelets in complement attack causes atypical hemolytic uremic syndrome. Blood 127, 2701–2710; 10.1182/blood-2015-11-680009 (2016).

14. Abeln, M. et al. Sialylation Is Dispensable for Early Murine Embryonic Development in Vitro. Chembiochem : a European journal of chemical biology 18, 1305–1316; 10.1002/cbic.201700083 (2017).

15. Rillahan, C. D. et al. Global metabolic inhibitors of sialyl- and fucosyltransferases remodel the glycome. Nature chemical biology 8, 661–668; 10.1038/nchembio.999 (2012).

16. Saqr, H. E., Pearl, D. K. & Yates, A. J. A review and predictive models of ganglioside uptake by biological membranes. Journal of neurochemistry 61, 395–411; 10.1111/j.1471-4159.1993.tb02140.x (1993).

17. Schneider, J. S., Sendek, S., Daskalakis, C. & Cambi, F. GM1 ganglioside in Parkinson’s disease: Results of a five year open study. Journal of the neurological sciences 292, 45–51; 10.1016/j.jns.2010.02.009 (2010).

18. Michalek, M. T., Bremer, E. G. & Mold, C. Effect of gangliosides on activation of the alternative pathway of human complement. Journal of immunology (Baltimore, Md. : 1950) 140, 1581–1587 (1988).

19. Horikawa, K. et al. Hemolysis of human erythrocytes is a new bioactivity of gangliosides. The Journal of experimental medicine 174, 1385–1391; 10.1084/jem.174.6.1385 (1991).

20. Pangburn, M. K. & Müller-Eberhard, H.J. Complement C3 convertase: cell surface restriction of beta1H control and generation of restriction on neuraminidase-treated cells. Proceedings of the National Academy of Sciences of the United States of America 75, 2416–2420; 10.1073/pnas.75.5.2416 (1978).

21. Lee, H., An, H. J., Lerno, L. A., German, J. B. & Lebrilla, C. B. Rapid Profiling of Bovine and Human Milk Gangliosides by Matrix-Assisted Laser Desorption/Ionization Fourier Transform Ion Cyclotron Resonance Mass Spectrometry. International Journal of Mass Spectrometry 305, 138–150; 10.1016/j.ijms.2010.10.020 (2011).

22. Schmidt, C. Q., Lambris, J. D. & Ricklin, D. Protection of host cells by complement regulators. Immunological reviews 274, 152–171; 10.1111/imr.12475 (2016).

23. Ladhani, S. N. et al. Invasive meningococcal disease in patients with complement deficiencies: a case series (2008-2017). BMC infectious diseases 19, 522; 10.1186/s12879-019-4146-5 (2019).

24. Mastellos, D. C. et al. Compstatin: a C3-targeted complement inhibitor reaching its prime for bedside intervention. European journal of clinical investigation 45, 423–440; 10.1111/eci.12419 (2015).

25. Souza, D. G., Esser, D., Bradford, R., Vieira, A. T. & Teixeira, M. M. APT070 (Mirococept), a membrane-localised complement inhibitor, inhibits inflammatory responses that follow intestinal ischaemia and reperfusion injury. British journal of pharmacology 145, 1027–1034; 10.1038/sj.bjp.0706286 (2005).

26. Blaum, B. S., Frank, M., Walker, R. C., Neu, U. & Stehle, T. Complement Factor H and Simian Virus 40 bind the GM1 ganglioside in distinct conformations. Glycobiology 26, 532– 539; 10.1093/glycob/cwv170 (2016).

27. Simons, M. et al. Exogenous administration of gangliosides displaces GPI-anchored proteins from lipid microdomains in living cells. Molecular biology of the cell 10, 3187–3196; 10.1091/mbc.10.10.3187 (1999).

28. The SASS Investigators. Ganglioside GM1 in acute ischemic stroke. The SASS Trial. Stroke 25, 1141–1148; 10.1161/01.str.25.6.1141 (1994).

29. Schnaar, R. L., Gerardy-Schahn, R. & Hildebrandt, H. Sialic acids in the brain: gangliosides and polysialic acid in nervous system development, stability, disease, and regeneration. Physiological reviews 94, 461–518; 10.1152/physrev.00033.2013 (2014).

30. Dalakas, M. C., Alexopoulos, H. & Spaeth, P. J. Complement in neurological disorders and emerging complement-targeted therapeutics. Nature reviews. Neurology 16, 601–617; 10.1038/s41582-020-0400-0 (2020).

31. Wada, T. & Nangaku, M. Novel roles of complement in renal diseases and their therapeutic consequences. Kidney international 84, 441–450; 10.1038/ki.2013.134 (2013).

32. Zipfel, P. F. & Skerka, C. Complement factor H and related proteins: an expanding family of complement-regulatory proteins? Immunology today 15, 121–126; 10.1016/0167-5699(94)90155-4 (1994).

33. Varki, A. et al. Symbol Nomenclature for Graphical Representations of Glycans. Glycobiology 25, 1323–1324; 10.1093/glycob/cwv091 (2015).

