## Supplement for "The monosialoganglioside GM1a protects trophoblasts, erythrocytes and endothelial cells against complement attack"

### Supplemental Methods

**Stem Cell Culture.** Basal TSC medium consisted of RPMI 1640 medium (biowest) with 1 % penicillin/streptomycin (PAN-biotech), 1mM sodium pyruvate (Merck), 50 µM 2-mercaptoethanol (Sigma-Aldrich) and 20 % fetal bovine serum (Merck). For stem cell culture, fetal bovine serum was heat inactivated for 30 min at 56 °C. Undifferentiated TSC were cultured with 70 % of the medium pre-conditioned on irradiated murine embryonic fibroblasts (Merck) supplemented directly before use with 1 µg/ml heparin (Sigma-Aldrich) and 25 ng/ml FGF4 (Reliatech) at 37°C in humidified incubators containing 5 % $CO_2$. Passaging of subconfluent cells was carried out using TrypLE (gibco). Medium was changed every other day. For differentiation experiments, standard differentiation medium (TSC medium without FGF4, Heparin and MEF conditioned medium) was used.

**xCGE-LIF.** For GSL analysis, glycan head groups of previously extracted glycosphingolipids were released by LudgerZyme Ceramide Glycanase (Ludger, Oxfordshire, UK), fluorescently labelled with 8-Aminopyrene-1,3,6-trisulfonic acid trisodium salt (APTS), and subjected to xCGE-LIF. For inter-sample quantitative comparison of signal intensities, a defined amount of APTS-labelled Glyko® Oligomannose 6 (Man6, Prozyme, Hayward, CA) was spiked into each sample. The obtained Man6 signal intensity was set to 1 normalized signal intensity (nRFU) and was used for normalization of peak intensities.

**Hemolysis assay.** Human whole blood was obtained by venipuncture and erythrocytes were isolated by centrifugation for 5 min at 1.000 x g. RBCs were washed two times in PBS and stored in Alsever's solution at 4°C with light agitation. Desialylation was performed under continuous agitation with 0.13 mg/ml purified *Arthrobacter ureafaciens* neuraminidase (AU 54)[1] for 30 min at 37°C in PBS. Simultaneously, sensitising of erythrocytes was achieved by incubation with anti-CD59 antibody (2 µg/ml). For the incorporation of GM1 or GM3, erythrocytes were incubated for 1 h at 37°C in PBS in agitation with the respective concentrations. Erythrocytes were washed and resuspended in DGHB-Mg-EGTA (HEPES 4.2 mM, NaCl 59 mM, $MgCl_2$ 7 mM, EGTA 10 mM, Glucose 2.08 % (w/v), Gelatine 0.08 % (w/v))[2] buffer to a final concentration of $5*10^7$ erythrocytes/ml. In a 96-well microtiter plate 50 µl erythrocytes were mixed with human serum diluted in DGHB Mg-EGTA to a final volume of 100 µl and incubated for 30 min at 37°C with continuous shaking. Subsequently, 150 µl NaCl 0.9% were added and the plate was centrifuged at 1000 xg for 5 min. Supernatant was transferred to a flat-bottomed 96-well plate and absorption at 414 nm was detected in a plate reader. All samples were measured in duplicates. Negative controls contain no serum. 100 % erythrocyte lysis was carried out by adding 200µl RBC lysis buffer ($NH_4Cl$ 155 mM, $NaHCO_3$ 12 mM, 0.1 mM EDTA) to 50 µl RBC. Experiments with human samples were approved by the local ethics committee (9371_BO_S_2020). Hemolytic assay with sheep red blood cells (SRBC) (Innovative research, Novi, MI, USA) were performed as described for human RBC. However, inucubation with anti-CD59 was omitted.

**Calcein release assay.** In brief, EA.hy926 cells ($10^4$ per well) were treated for 30 minutes at 37°C with Neu, incubated for 30 minutes with Calcein AM (25 µM) and GM1a or GM3 (Avanti Polar Lipids, United States) of varying concentrations was added for 1h at 37°C. To induce a complement attack the cells were then incubated with 15% human serum together with anti-CD59 (10 µg/ml). After 30 minutes at 37°C fluorescence of the supernatant at 494 nm was measured.

**Supplemental Figures**

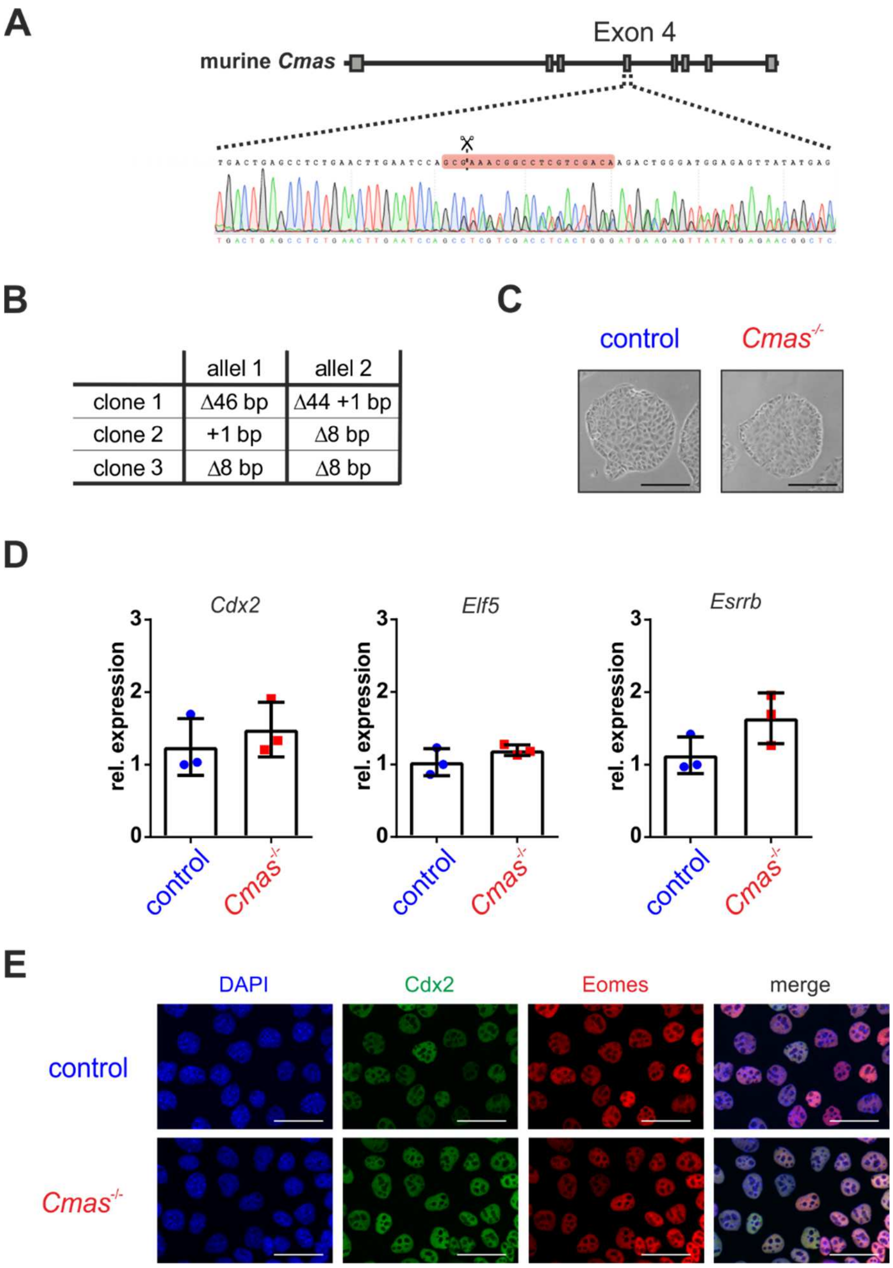


**Figure S1. Genetic deletion of *Cmas* does not alter the stem cell status of TSC.** (**A**) Scheme of the murine Cmas gene and sequencing data of one representative *Cmas*$^{-/-}$ clone with indels at the target site in exon 4. (**B**) The table depicts the size of indels in exon 4 of *Cmas*$^{-/-}$ clones determined by sequencing. (**C**) Brightfield images of control and one representative *Cmas*$^{-/-}$ TSC cultivated under standard conditions. Scale bars = 250 µm. (**D**) Quantitative PCR of the TSC marker *Cdx2*, *Elf5* and *Esrrb* from control and one representative *Cmas*$^{-/-}$ TSC clone. Gene expression was plotted relative to control TSC. n=3 Mean values are depicted with standard deviation. (**E**) Detection of EOMES and CDX2 by indirect immunofluorescence analysis of control and one representative *Cmas*$^{-/-}$ TSC clone. Nuclei were stained with DAPI and are shown in blue. Scale bars = 50 µm.

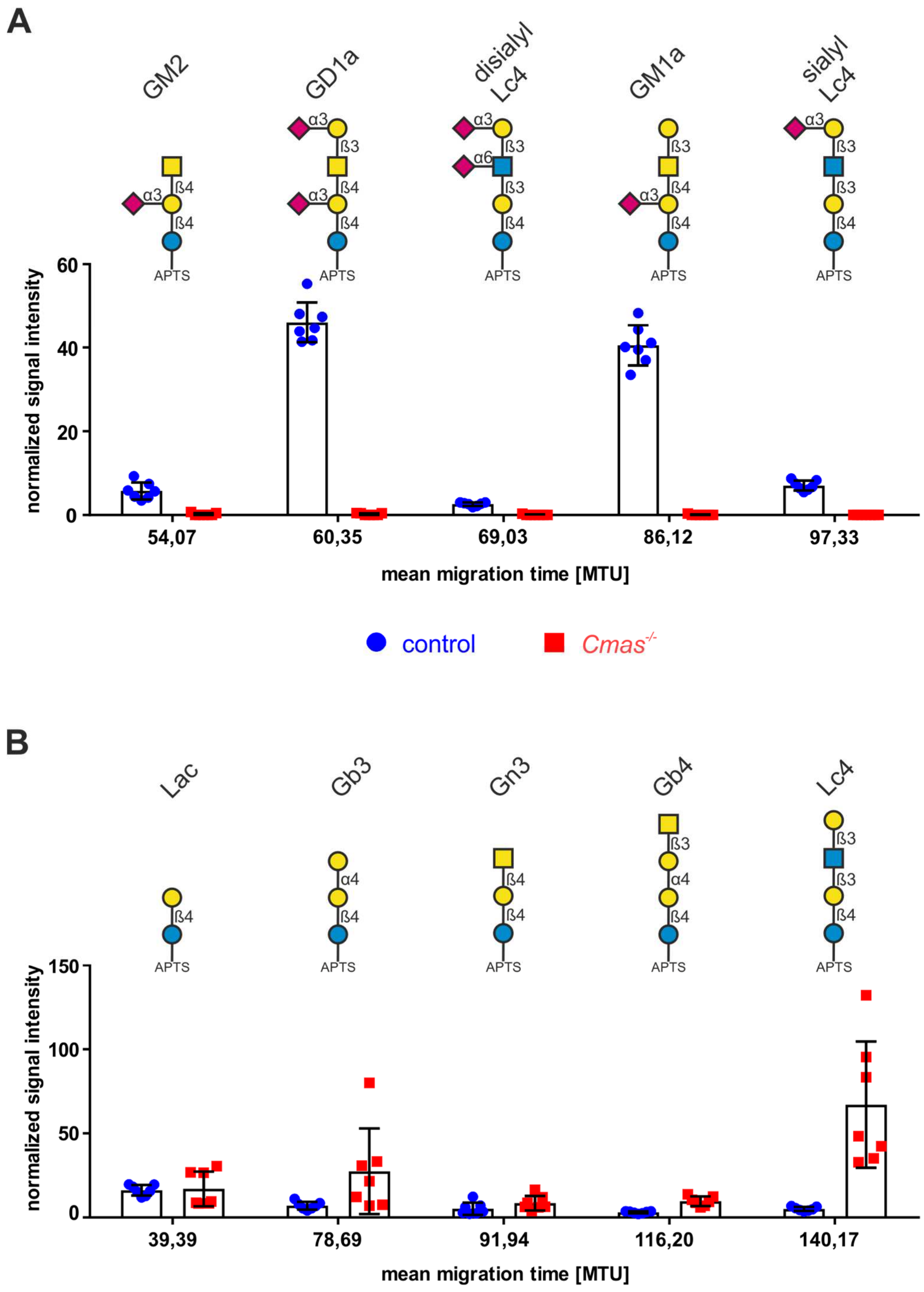


**Figure S2. *Cmas*$^{-/-}$ TSC lack sialylated glycosphingolipids.** Quantification of sialylated (**A**) and neutral (**B**) glycans derived from glycosphingolipids of control and one representative *Cmas*$^{-/-}$ TSC clone using xCGE-LIF. Glycan notations for GM2, GD1a and GM1a refer to the respective glycosphingolipid derived glycan. For inter-sample comparisons signal intensities were normalized to Man6 (nRFU). MTU: migration time unit. Symbol nomenclature according to [3]. n=7 Mean values are depicted with standard deviation.

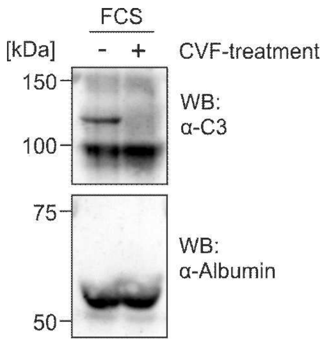


**Figure S3. Depletion of C3 in FCS by CVF analyzed by Western Blot.** Control and CVF treated FCS was separated by SDS-PAGE and immunostained with anti-C3 antibody. Anti-albumin staining was used as loading control.

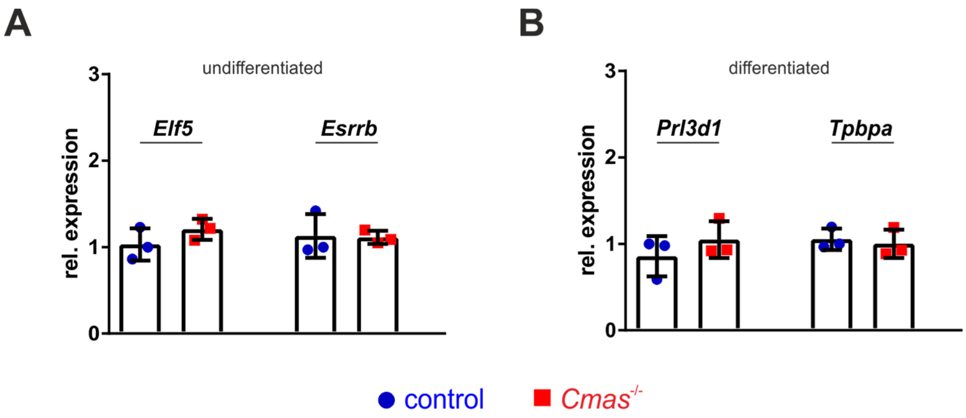


**Figure S4. Unaltered expression of TSC stemness and differentiation markers in *Cmas*$^{-/-}$ TSC under complement deprived conditions.** Quantitative PCR of TSC markers Elf5 and Esrrb and differentiation markers Prl3d1 and Tpbpa from control and one representative *Cmas*$^{-/-}$ TSC clone in undifferentiated and 7 day differentiated cells in medium with heat inactivated FCS (HI-FCS). Gene expression was plotted relative to control TSC. n=3. Mean values are depicted with standard deviation.

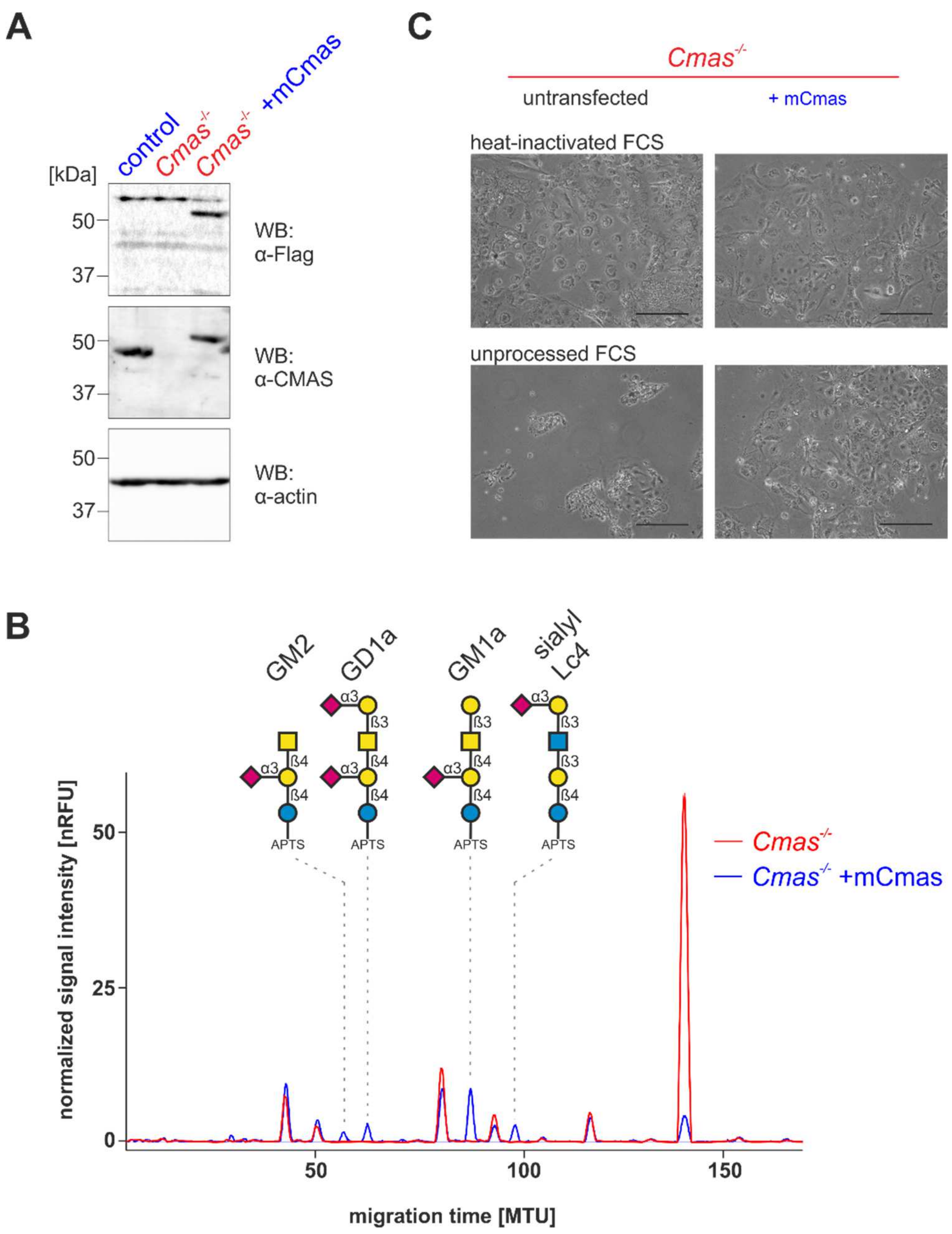


**Figure S5. Recomplementing of *Cmas*$^{-/-}$ TSC with murine Cmas rescues *Cmas*$^{-/-}$ TSC from complement attack.** (**A**) Control, *Cmas*$^{-/-}$ and recomplemented *Cmas*$^{-/-}$ TSC were lysed and separated by SDS-PAGE, blotted onto PVDF membrane and immunostained with anti-CMAS antibody and anti-Flag antibody. Anti-actin immunostaining was used as loading control. (**B**) Overlay of xCGE-LIF electropherograms of APTS-labeled GSL-derived glycans of *Cmas*$^{-/-}$ and recomplemented *Cmas*$^{-/-}$ TSC. Glycan notations for GM2, GD1a and GM1a refer to the respective glycosphingolipid derived glycan. For inter-sample comparisons signal intensities were normalized to Man6 (nRFU). MTU: migration time unit, n=3. Symbol nomenclature according to [3]. (**C**) Brightfield images of *Cmas*$^{-/-}$ and recomplemented *Cmas*$^{-/-}$ TSC after 7 days of differentiation in medium with unprocessed, heat-inactivated FCS. Scale bars = 250 µm.

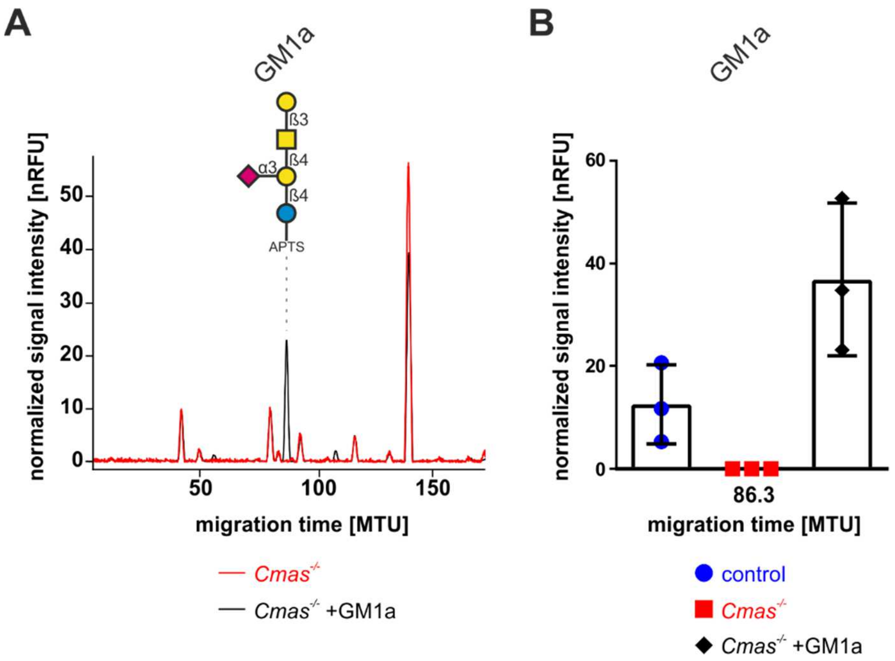


**Figure S6. TSC incorporate exogenously administered GM1a.** (**A**) Overlay of xCGE-LIF electropherograms of APTS-labeled GSL-derived glycans of *Cmas*$^{-/-}$ TSC with and without exogenously administered GM1a. Glycan notation for GM1a refers to the glycosphingolipid derived glycan. For inter-sample comparisons signal intensities were normalized to Man6 (nRFU). MTU: migration time unit, n=3. Symbol nomenclature according to [3]. (**B**) Quantification of GM1a in control and *Cmas*$^{-/-}$ TSC with and without exogenously administered GM1a, n=3. Mean values are depicted with standard deviation.

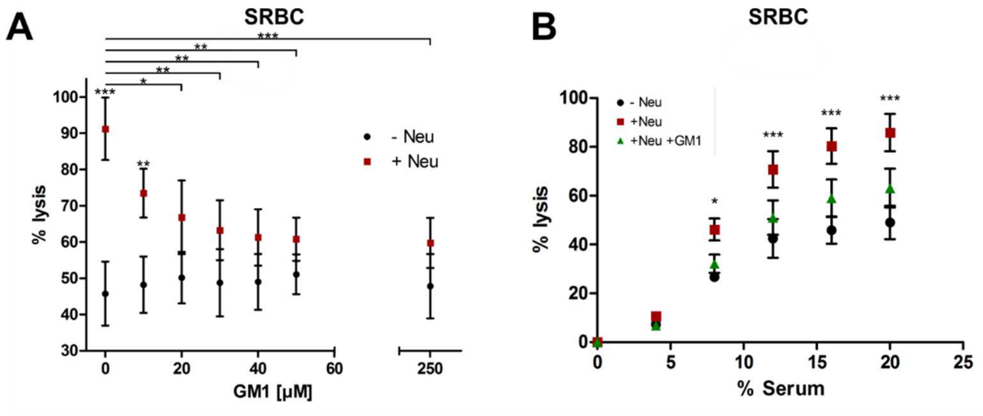


**Figure S7. GM1a restores complement resistance of sheep red blood cells.** (**A**) SRBC were left untreated or sensitised for activation of the alternative complement pathway by incubation with Neu. Furthermore, control and Neu treated SRBC were incubated with increasing concentrations of GM1a. Hemolysis was determined at 415 nm after incubation of SRBC with 16.6 % human complement active serum in DGHB-Mg-EGTA. Hemolysis was plotted relative to SRBC incubated with only red cell lysis buffer, which was defined as 100% lysis. n=3 (**B**) Control, Neu sensitised or Neu sensitised and subsequently GM1a (50 µM) treated SRBC were incubated with increasing concentrations of human complement active serum in DGHB-Mg-EGTA. Hemolysis was plotted relative to SRBC incubated with only red cell lysis buffer, which was defined as 100% lysis. n=3. Untreated SRBC showed 45.75 (± 5.09) % lysis whereas 91.23 (± 4.97) % of Neu sensitized SRBC were lysed. GM1a treatment of Neu sensitized SRBC significantly improved the resistance to complement dependent lysis of SRBC in a dose dependent manner, with maximal inhibition of lysis at 50 µM GM1a (SRBC 51.07 ± 3.16 %; Neu sensitized SRBC 60.77 ± 3.42 %). Additional increase of GM1a up to a concentration of 250 µM did not further improve Neu sensitized SRBC survival (SRBC 47.80 ± 5.13 %; Neu sensitized SRBC 59.76 ± 3.99 %). Thus, administration of 50 µM GM1a restored complement resistance to 84% of native SRBC. The same effect was observed when Neu sensitized SRBC, with and without subsequent treatment with 50 µM GM1a, were challenged with increasing concentrations of human serum. For statistical analysis a two-way ANOVA with Bonferroni's multiple comparison post-test was performed * P < 0.05, ** P < 0.01, *** P < 0.001. Mean values are depicted with standard deviation.

**Supplemental Tables**

**Table S1: qPCR primer sequences**

| | |
|---|---|
| *Cdx2 fwd* | TCCTGCTGACTGCTTTCTGA |
| *Cdx2 rev* | CCCTTCCTGATTTGTGGAGA |
| *Elf5 fwd* | ATTCGCTCGCAAGGTTACTCC |
| *Elf5 rev* | GGATGCCACAGTTCTCTTCAGG |
| *Esrrb fwd* | AGTACAAGCGACGGCTGG |
| *Esrrb rev* | CCTAGTAGATTCGAGACGATCTTAGTCA |
| *Pgk1 fwd* | CTGACTTTGGACAAGCTGGACG |
| *Pgk1 rev* | GCAGCCTTGATCCTTTGGTTG |
| *Prl3d1 fwd* | TGGAGCCTACATTGTGGTGG |
| *Prl3d1 rev* | TGGCAGTTGGTTTGGAGGA |
| *Sdha fwd* | TGGTGAGAACAAGAAGGCATCA |
| *Sdha rev* | CGCCTACAACCACAGCATCA |
| *Tpbpa fwd* | CCAGCACAGCTTTGGACATCA |
| *Tpbpa rev* | AGCATCCAACTGCGCTTCA |
| *Ywhaz fwd* | TTGATCCCCAATGCTTCGC |
| *Ywhaz rev* | CAGCAACCTCGGCCAAGTAA |

**Table S2: Antibodies and lectins**

| Antibody | Source | Label | Manufacturer | Catalog no. | Dilution |
|---|---|---|---|---|---|
| Anti-Albumin | goat | - | Abcam | ab19194 | WB: 1:5000 |
| Anti-Actin | mouse | - | Millipore | #MAB1501 | WB: 1:100000 |
| Anti-Cdx2 | rat | - | Abcam | ab76541 | IF: 1:300 |
| Anti-CD59 | rat | - | BioRad | MCA715G | HA: 2 µg/ml<br>CRA: 10 µg/ml |
| Anti-Cmas serum | rabbit | - | made in-house [4] | - | WB: 1:15000 |
| Anti-C3 | goat | - | MP Biomedicals | #55463 | WB: 1:5000<br>IF: 1:200 |
| Anti-Eomes | rabbit | - | Invitrogen | 14-4875-82 | IF: 1:100 |
| Anti-Factor H | goat | - | Complement Technology | A237 | IF: 1:1000 |
| Anti-Flag | rabbit | - | Cell Signaling | 2368 | WB: 1:5000 |
| Cholera Toxin B Subunit | vibrio cholera | FITC | Sigma-Aldrich | C9903 | IF: 1:200 |
| Anti-rat-IgG | goat | Alexa488 | Invitrogen | A11006 | IF: 1:500 |
| Anti-rabbit-IgG | sheep | Cy3 | Sigma-Aldrich | C2306 | IF: 1:500 |
| Anti-goat-IgG | donkey | Alexa488 | Invitrogen | A11055 | IF: 1:500 |
| Anti-goat-IgG | rabbit | HRP | Jackson | 305-035-003 | WB: 1:15000 |
| Anti-rabbit-IgG | goat | HRP | Jackson | 111-035-003 | WB: 1:15000 |
| Anti-mouse-IgG+IgA+IgM | goat | HRP | Southern Biotech | 1010-05 | WB: 1:15000 |

### References


1. Christensen, S. & Egebjerg, J. Cloning, expression and characterization of a sialidase gene from Arthrobacter ureafaciens. *Biotechnology and applied biochemistry* **41,** 225–231; 10.1042/BA20040144 (2005).
2. Moreno-Indias, I., Dodds, A. W., Argüello, A., Castro, N. & Sim, R. B. The complement system of the goat: haemolytic assays and isolation of major proteins. *BMC veterinary research* **8,** 91; 10.1186/1746-6148-8-91 (2012).
3. Varki, A. *et al.* Symbol Nomenclature for Graphical Representations of Glycans. *Glycobiology* **25,** 1323–1324; 10.1093/glycob/cwv091 (2015).
4. Schaper, W. *et al.* Identification and biochemical characterization of two functional CMP-sialic acid synthetases in Danio rerio. *The Journal of biological chemistry* **287,** 13239–13248; 10.1074/jbc.M111.327544 (2012).


**Full uncropped gels**

**Figure 1, B**

**Anti CMAS Western Blot**

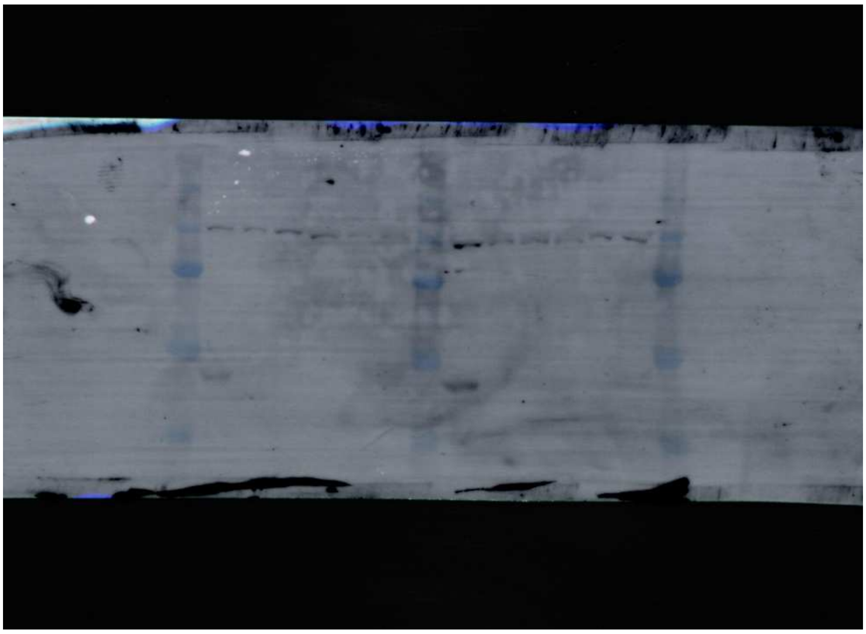

**Anti Actin**

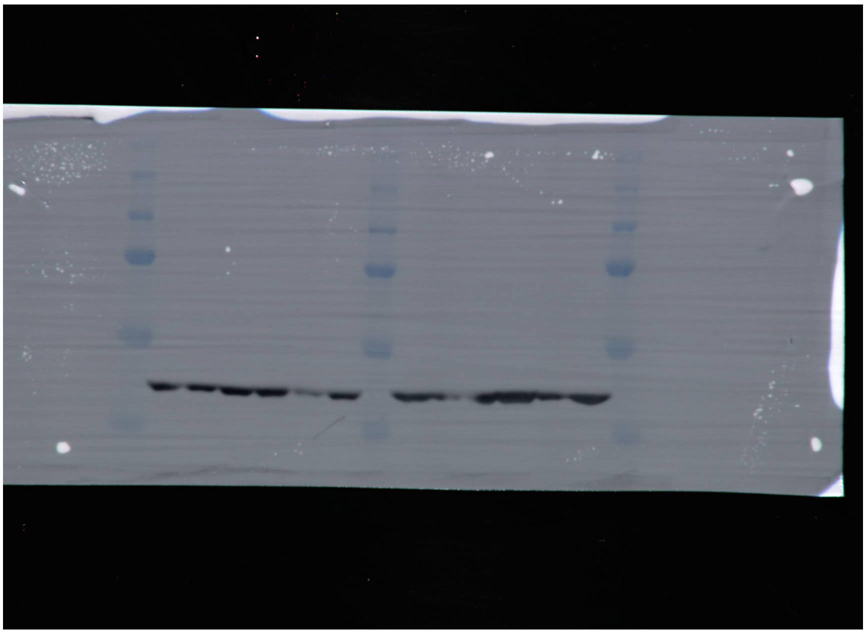

**Figure S3**

**Anti C3** **Anti Albumin**

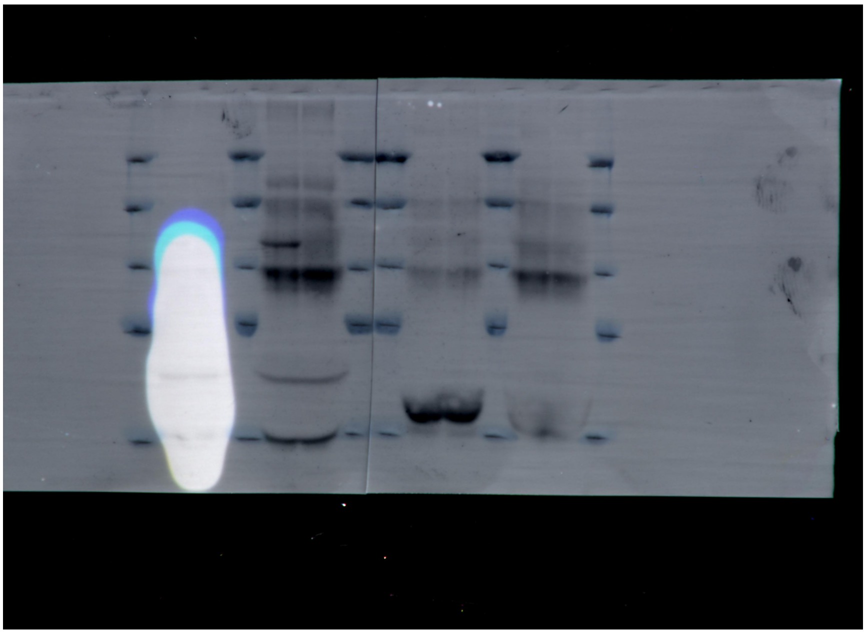

**Figure S5, A**

**Anti Flag**

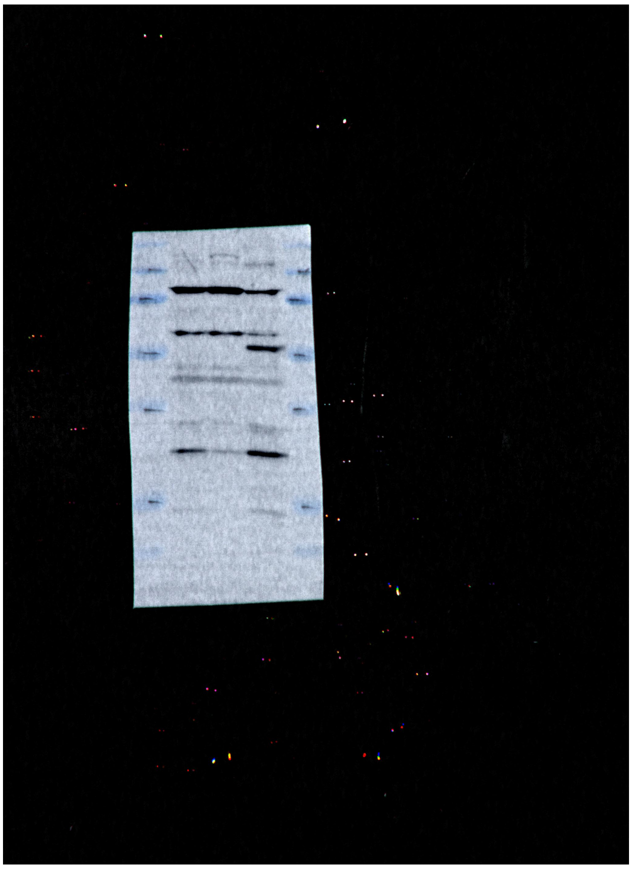

**Anti Actin** **Anti CMAS**

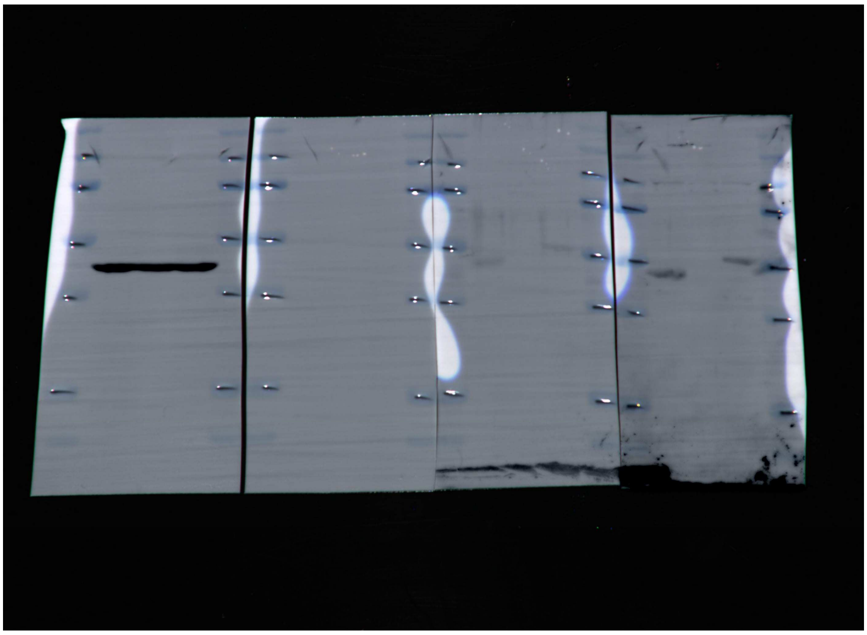
